# Selective observation of semi-rigid non-core residues in dynamically complex mutant huntingtin protein fibrils

**DOI:** 10.1101/2022.05.13.489937

**Authors:** Irina Matlahov, Jennifer C. Boatz, Patrick C. A. van der Wel

**Affiliations:** Zernike Institute for Advanced Materials, University of Groningen, Groningen, The Netherlands; Department of Structural Biology, University of Pittsburgh School of Medicine, Pittsburgh, PA, USA

**Keywords:** intermediate-timescale motion, solid state NMR, Huntington’s disease, polyglutamine, dynamics

## Abstract

Many amyloid-forming proteins, which are normally intrinsically disordered, undergo a disorder-to-order transition to form fibrils with a rigid β-sheet core flanked by disordered domains. Solid-state NMR (ssNMR) and cryogenic electron microscopy (cryoEM) excel at resolving the rigid structures within amyloid cores but studying the dynamically disordered domains remains challenging. This challenge is exemplified by mutant huntingtin exon 1 (HttEx1), which self-assembles into pathogenic neuronal inclusions in Huntington disease (HD). The mutant protein’s expanded polyglutamine (polyQ) segment forms a fibril core that is rigid and sequestered from the solvent. Beyond the core, solvent-exposed surface residues mediate biological interactions and other properties of fibril polymorphs. Here we deploy magic angle spinning ssNMR experiments to probe for semi-rigid residues proximal to the fibril core and examine how solvent dynamics impact the fibrils’ segmental dynamics. Dynamic spectral editing (DYSE) 2D ssNMR based on a combination of cross-polarization (CP) ssNMR with selective dipolar dephasing reveals the weak signals of solvent-mobilized glutamine residues, while suppressing the normally strong background of rigid core signals. This type of ‘intermediate motion selection’ (IMS) experiment based on cross-polarization (CP) ssNMR, is complementary to INEPT- and CP-based measurements that highlight highly flexible or highly rigid protein segments, respectively. Integration of the IMS-DYSE element in standard CP-based ssNMR experiments permits the observation of semi-rigid residues in a variety of contexts, including in membrane proteins and protein complexes. We discuss the relevance of semi-rigid solvent-facing residues outside the fibril core to the latter’s detection with specific dyes and positron emission tomography tracers.

**Highlights:**

- Mutant huntingtin exon 1 fibrils feature a broad range of molecular dynamics.
- Molecular motion is coupled to water dynamics outside the fiber core.
- Dynamics-based spectral editing ssNMR reveals mobile non-core residues.
- Intermediate-motion selection via dipolar dephasing of rigid sites.
- Semi-mobile glutamines outside the fiber core observed and identified.

## Introduction

Self-assembled protein filaments in the form of amyloids or amyloid-like structures arise in many places in the human body. Pathogenic amyloids are culprits of neurodegenerative aggregation diseases like Alzheimer’s disease (AD), Parkinson’s disease (PD), and Huntington’s disease (HD) (Chiti and Dobson, 2017). Structural studies of these types of fibrillar assemblies have seen great progress enabled by both solid-state NMR spectroscopy (ssNMR) and cryogenic electron microscopy (cryoEM) (Fitzpatrick et al., 2017; Fitzpatrick and Saibil, 2019; Hervas et al., 2020; Lutter et al., 2021; van der Wel, 2017). This has revealed that these fibrils feature a variety of β-sheet based fibril cores, which are internally remarkably well ordered, even when a single protein can often form distinct fibril strains (or polymorphs).

Within polymorphic fiber architectures, the focus is typically on differences in the structure of the cross-β fiber core. However, segments or domains outside this core region play a critical role in the misfolding process and help determine the functional (and toxic) properties of the aggregates. ‘Flanking’ segments do not partake in the β-sheet fiber core and are surface exposed, where they mediate interactions with other cellular components (Lin et al., 2017; Ulamec et al., 2020). The water-facing interfaces of the β-sheet fiber core itself are of interest in a diagnostic context, as they are targets for amyloid-specific fluorescent dyes and tracers used in positron emission tomography (PET) (Fändrich et al., 2018). Thus, a detailed understanding of the residues and segments outside the rigid fibril core is of substantial biomedical interest.

Mutant huntingtin (Htt) fragments implicated in Huntington’s disease (HD) generate pathogenic protein aggregates (Bates et al., 2015). In this inherited neurodegenerative disease, N-terminal fragments of mutant huntingtin (Htt) form ordered protein aggregates, which are found as cytoplasmic and nuclear inclusions in patient neurons (Macdonald, 1993). Among these fragments, much interest is in the first exon of Htt (HttEx1) which bears the HD mutation, induces HD-like symptoms in model animals and has been detected in patient-derived inclusions. Notably, the mutation behind HD involves the genetic expansion of the HttEx1 polyglutamine (polyQ) domain, as illustrated in Figure 1 (Scherzinger et al., 1997). This domain is flanked by an N-terminal Htt^NT^ segment and a C-terminal proline-rich domain (PRD).

**Figure 1.**
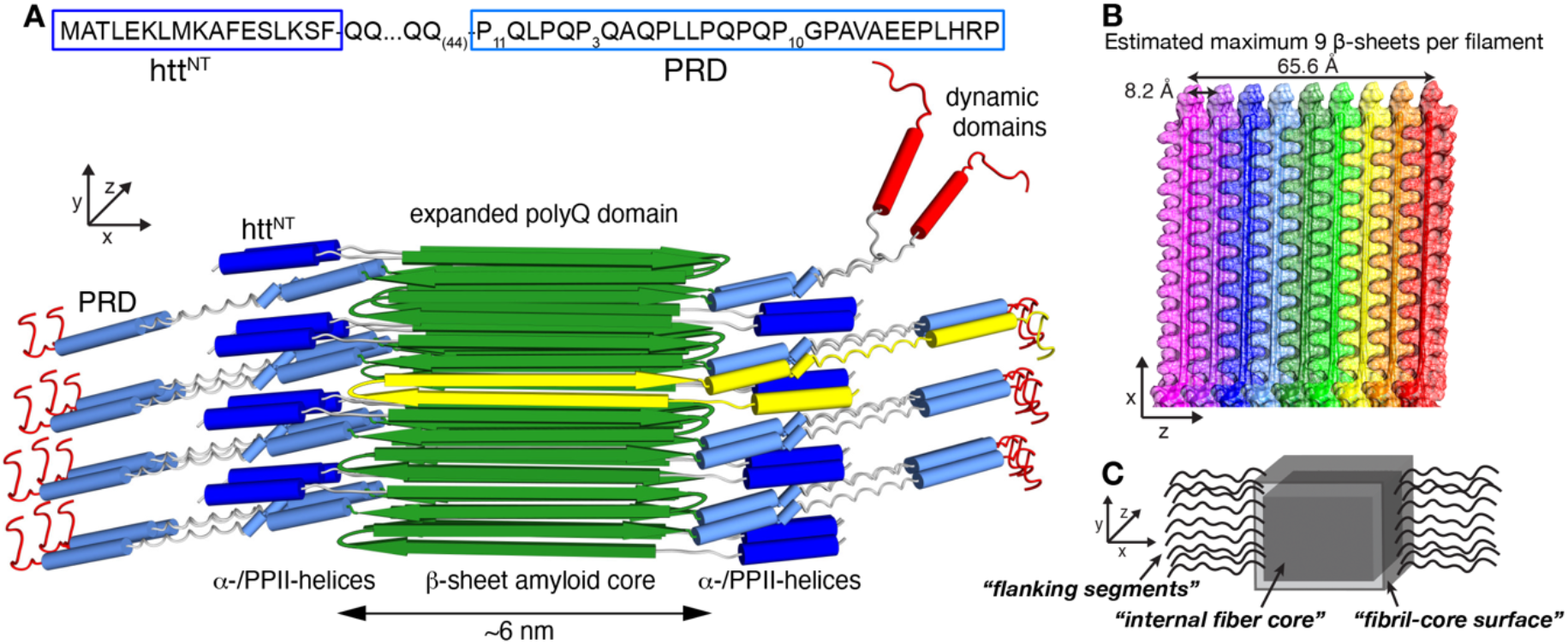
Fibril architecture of aggregated mutant HttEx1. (A) Sequence and structural model of HttEx1 fibers. The xyz axes indicate the fiber growth direction (y), fiber width (x) and fiber depth (z). The yellow β-hairpin structure shows a monomer, which is exposed one the front side of the fibril core. (B) Structural model of HttEx1 fibers showing the layering of β-sheets in the z direction. Each β-sheet is shown in different colour, with a 6-7 nm protofilament core composed of around nine stacked β-sheets. (C) Schematic drawing of the fibril structure, marking the nomenclature distinguishing the internal fiber core (i.e. dry interfaces), fibril core surface (Eisenberg “wet interface”), as well as the solvent exposed flanking domains. Panels A-B adapted with permission from ref. (Lin et al., 2017) and (Boatz et al., 2020), under the Creative Commons CC-BY license.

The structure of polyQ-expanded HttEx1 aggregates has been studied by many techniques (Bugg et al., 2012; Duim et al., 2014; Hoop et al., 2014; Bäuerlein et al., 2017; Caulkins et al., 2018; Guo et al., 2018; Boatz et al., 2020; Galaz-Montoya et al., 2021; Isas et al., 2021; Nazarov et al., 2022). Although an atomic level structure is not yet available, key features of the HttEx1 fibrils are known (Figure 1). Their stable core is formed by the polyQ domain, forming a rigid, multilayered β-sheet structure that is internally devoid of solvent molecules (Sharma et al., 2005; Schneider et al., 2011; Hoop et al., 2016; Matlahov and van der Wel, 2019; Nazarov et al., 2022). Here, we refer to the rigid and solvent-sequestered (“dehydrated”) polyQ residues as the “internal fiber core” (Figure 1C). Outside this internal core, there are two types of exposed features: polyQ residues that face the solvent as well as non-polyQ flexible flanking segments. Figure 1A shows a schematic drawing, in which the flanking segments are shown: the N-terminal (htt^NT^) (Figure 1A, dark blue cylinders) and C-terminal PRD (light blue cylinders). These segments are distinct from the polyQ core based on higher mobility and solvent exposure (Bugg et al., 2012; Lin et al., 2017) (Figure 1A). The htt^NT^ may adopt a dynamic α-helical structure or disordered and flexible state, and is characterized by a significant proportion of hydrophobic and methyl-bearing residues (Table 1) (Boatz et al., 2020; Isas et al., 2015). The PRD domain contains two PPII helical structures with varying levels of mobility (Bugg et al., 2012; Caulkins et al., 2018; Lin et al., 2017). The layered polyQ core structure results in a fibril geometry in which the rigid core is on two sides bounded by the dynamic flanking domains, and on two other sides presenting outward-facing glutamines. The latter sides are reminiscent of the ‘wet’ interface of amyloidogenic steric zipper peptides previously studied by the Eisenberg group (Nelson et al., 2005). This wet interface we will call the “fibril-core surface” (see Figure 1C). These water-facing residues are expected to display semi-rigid behavior, being on the one hand exposed to mobile water while also anchored in the immobilized β-sheet structure of the amyloid-like core.

**Table 1.** Distribution of amino acids between the domains of the Q44-HttEx1 protein.

| Domain | Q | P | L | H | A | E | S | K | G | M | F | T | V | R |
| --- | --- | --- | --- | --- | --- | --- | --- | --- | --- | --- | --- | --- | --- | --- |
| Htt<sup>NT</sup> | - | - | 3 | - | 2 | 2 | 2 | 3 | - | 2 | 2 | 1 | - | - |
| PolyQ | 44 | - | - | - | - | - | - | - | - | - | - | - | - | - |
| PRD | 6 | 31 | 4 | 1 | 3 | 2 | - | - | 1 | - | - | - | 1 | 1 |
| His-tag | - | - | - | 6 | - | - | 2 | - | 1 | - | - | - | - | - |
| <u>Total:</u> | 50 | 31 | 7 | 7 | 5 | 4 | 4 | 3 | 2 | 2 | 2 | 1 | 1 | 1 |

Our current understanding of the abovementioned fibril structure stems from an integration of structural biology techniques, with a prominent role for ssNMR spectroscopy. Of particular importance have been ssNMR experiments that distinguish the rigid core from the flexible flanking segments, enabled by dynamic spectral editing (DYSE) techniques (Matlahov and Van der Wel, 2018). With some exceptions (Kashefi et al., 2019), DYSE methods usually are focused on revealing either highly mobile residues or the most rigid parts of the sample and have proved effective for many different protein assemblies (Helmus et al., 2008; Murray et al., 2017; van der Wel, 2017).

Yet, it has remained challenging to observe and study residues or segments with an intermediate mobility. Some partly immobilized domains (that are not fully rigid) can be detected by CP-based ssNMR (Helmus et al., 2010; Ryder et al., 2021), although at reduced intensity. The signals from such semi-mobile domains are often masked by stronger signals from more rigid parts of the protein structure. In HttEx1 fibrils studied by CP ssNMR, most of the polyQ ssNMR signal reflects residues internal to the rigid core (see also Boatz et al. 2020). Yet, there is also a direct polyQ-core-solvent interface (Figure 1B,C, “fibril-core surface”) that is of real interest, as it harbors binding sites for amyloid-specific dyes (like congo red or thioflavin T) and PET ligands designed for *in vivo* imaging of mutant Htt (Liu et al., 2020; Bertoglio et al., 2021, 2022). Moreover, the fibril-core surface may be implicated in secondary nucleation events that play a prominent role in aggregation of HttEx1 and other amyloidogenic polypeptides (Boatz et al., 2020; Wagner et al., 2018).

In this work, we set out to gain more insight into the functionally relevant, but difficult-to-observe segments and residues outside the HttEx1 fiber core. We deploy variable temperature ssNMR studies to probe the role of solvent dynamics in modulating the mobility of the fibrils, juxtaposing the buried fiber core and exposed domains. Moreover, we deploy a new DYSE method that combines two previously reported techniques (^1^H-^13^C DIPolar chemical SHIFT (DIPSHIFT) and ^13^C-^13^C Dipolar Assisted Rotational Resonance (DARR) experiments (Munowitz et al., 1981; Takegoshi et al., 2001)), to selectively probe the semi-mobile residues such as those that are exposed but proximal to the rigid core. This approach aims to reduce the disadvantageous dynamic noise problem affecting the detection of semi-mobile surface glutamines by partly or fully suppressing the signals from rigid polyQ core residues, complementing a recently reported technique based on other types of dipolar interactions (Kashefi et al., 2019). With this technique we can identify otherwise invisible signals of semi-rigid glutamine residues that are distinct from the dominant signals from the rigid core.

## Materials and Methods

### Protein expression, purification, and fibrillation

Mutant huntingtin exon 1 with a 44-residue polyQ core was expressed as part of a maltose binding protein (MBP) fusion protein as previously reported (Hoop et al., 2014; Lin et al., 2017). The fusion protein MBP-Q44-HttEx1 was subcloned into a pMalc2x plasmid by Genscript. The protein expression was done in *Escherichia coli* BL21(DE3) pLysS cells (Invitrogen, Grand Island, NY). Samples for MAS ssNMR were uniformly ^13^C, ^15^N labeled with ^13^C D-glucose and ^15^N ammonium chloride. After protein expression, cells were pelleted at 7000g, resuspended in phosphate buffered saline (PBS) (SKU BP399-4, Fisher BioReagents, Pittsburgh, PA) and lysed in presence of 1 mM phenylmethanesulfonyl fluoride (PMSF) (SKU 215740050, Acros Organics, New Jersey, USA) by an M-110Y high pressure pneumatic high shear fluid processor (Microfluidics, Newton, MA). Subsequently, cells were centrifuged at 38,720g for 1h using a Sorvall RC 3C Plus centrifuge (Thermo Scientific, Waltham, MA). The supernatant that contained the soluble fusion protein was filtered over Millex-GP syringe-driven 0.22-µm PES membranes (Millipore Sigma, Burlington, MD). The fusion protein was purified by fast protein liquid chromatography (FPLC) using a 5ml HisTrap HP nickel column (GE Healthcare, Uppsala, Sweden) with 0.5M imidazole gradient (SKU I5513-100G, Sigma, St. Louis, MO) on an AKTA system (GE Healthcare, Chicago, IL). The imidazole was removed from the purified protein using an Amicon Ultra centrifugal filter with a regenerated cellulose membrane (Millipore Sigma, Burlington, MA). At least 3 washes with imidazole-free PBS buffer were done. Protein concentration was calculated from the absorbance at 280nm. According to ProtParam tool by ExPasy (Gasteiger, 2003) the extinction coefficient of the fusion protein is 66,350 M^-1^cm^-1^. Factor Xa protease (SKU PR-V5581, Promega, Madison, WI) was used to cleave MBP-Q44-HttEx1 at 22 °C (Boatz et al., 2020; Lin et al., 2017) to release Q44-HttEx1. After 3 days the obtained fibrils were washed with PBS to remove the cleaved MBP.

### NMR experiments

The hydrated U-^13^C,^15^N-labeled Q44-HttEx1 fibrils were packed into a 3.2mm ssNMR rotor (Bruker Biospin) by sedimentation, using a previously described ultracentrifugal packing tool (Mandal et al., 2017). Experiments were performed on a wide-bore Bruker Avance-I 600 MHz spectrometer at 277K, 270K, 263K and 256K temperatures using a triple-channel (HCN) 3.2mm MAS EFree probe. All ^13^C-detected experiments were done using two-pulse phase modulated (TPPM) proton decoupling of 83 kHz during acquisition (Bennett et al., 1995). Direct excitation ^1^H MAS NMR experiments were performed using a 3µs 90° pulse on ^1^H and recycle delay of 2.8s. 1D ^13^C direct excitation experiments were done using a 90° ^13^C pulse of 4µs and a 3s recycle delay, along with high power ^1^H decoupling during acquisition. This recycle delay is smaller than 5x ^13^C T_1_ of the polyQ core residues, resulting in a suppression of these core signals relative to more mobile residues (Sivanandam et al., 2011). 1D ^13^C cross-polarization (CP) experiments were done using a 3µs 90° pulse on ^1^H, a contact time of 1ms, recycle delay of 3s. 2D ^13^C-^13^C DARR experiments (Takegoshi et al., 2001) were performed using a 3µs 90° pulse on ^1^H, 4 µs 90° pulses on ^13^C, a ^1^H-^13^C CP contact time of 1ms or 750 µs at 277 and 256K respectively, a DARR mixing time of 25ms, and a recycle delay of 2.8s. A 2D PDSD (proton-driven spin diffusion) experiment was done at 256K using same pulses as in 2D DARR experiment but with a mixing time of 500ms. 2D ^13^C-^15^N NCO experiments were done using a 3µs 90° pulse on ^1^H, 8µs 180° pulse on ^13^C, ^1^H-^15^N contact time of 1.5ms (277K) and 800µs (256K), ^15^N-^13^C contact time of 4ms and recycle delay of 2.8s. 2D ^13^C-^15^N NCA experiments were done using a recycle delay of 2.8s, 3µs 90° pulse on ^1^H, 8µs 180° pulse on ^13^C, 1.5ms (277K) and 800µs (256K) and 4ms ^1^H-^15^N and ^15^N-^13^C contact times, respectively. In both NCA and NCO experiments, the power levels for ^15^N and ^13^C during N-C transfer steps were 50kHz and 62.5kHz, respectively. The pulse shapes for H-N and N-C magnetization transfer were ramp70100 (70-100% power ramp) and square.100 (no ramp) respectively. The 1D DIPSHIFT experiments (Munowitz et al., 1981) were performed according to previously reported procedures (Hoop et al., 2014; Lin et al., 2017). This involved the use of a R18_1_^7^ R-sequence (Zhao et al., 2001) for ^1^H-^13^C dipolar protein recoupling. The 2D DIPSHIFT-DARR experiments were performed using recycle delay of 2.8s, 3µs 90° pulse on ^1^H, 4 and 8µs 90° and 180° ^13^C pulses respectively, 750µs ^1^H-^13^C contact time, 25ms mixing time. The ssNMR measurements were performed using Bruker Topspin software. Spectral processing and analysis were done using Topspin, CcpNMR Analysis (Skinner et al., 2016), as well as MestreNova software. Peak integration and peak deconvolution in the 1D slices were done using MestReNova. The ^1^H, ^13^C, and ^15^N chemical shifts were referenced using indirect and external referencing by measuring the ^13^C signals of adamantane (Harris et al., 2008; Morcombe and Zilm, 2003). Briefly, the adamantane ^13^CH_2_ signal was assumed to have a chemical shift of 40.49 ppm relative to aqueous DSS, and referencing of other nuclei was done by using the relevant Ω values (^1^H referenced to aqueous DSS; ^15^N to liquid ammonia). Scripts that facilitate this process are available from the authors. Additional experimental details are tabulated in the Table S1 in the supporting information.

## Results

### Variable dynamics of distinct segments in the fibrils

We have previously described our protocols for generating amyloid-like fibrils from the mutant HttEx1 protein outfitted with a 44-residue polyQ segment (Q44-HttEx1) and uniformly labeled with ^13^C and ^15^N (Hoop et al., 2014; Lin et al., 2017). These ssNMR samples contain hydrated protein fibrils, with an excess of buffer that permits free motion of the solvent exposed segments such as semi-rigid parts of the polyQ domain (fibril-core surface), as well as N- and C-terminal flanking segments. First, we describe our use of variable temperature ssNMR studies to probe how solvent motion impacts the rigid and semi-rigid fiber components. **Figure 2** (top) shows the 1D ^13^C spectra at 277K, obtained with either CP-based ssNMR (**Figure 2**A) or direct excitation (**Figure 2**B), which are differently sensitive to mobility. Rigid sites dominate the CP spectra, with highly flexible residues being invisible. The direct excitation ssNMR shows rigid and mobile sites alike. Based on prior 2D ssNMR (Lin et al., 2017; Sivanandam et al., 2011) we can attribute assignments to the peaks (marked in **Figure 2**). The most rigid sites (high peaks in **Figure 2**A) are from the polyQ core which composed of two Gln conformers marked Qa and Qb. In contrast, increased mobility is seen for the protein’s other domains (htt^NT^ and PRD), which unlike the polyQ segment contain various other amino acid types (Table 1). Figure 2B shows that such residues (e.g. marked A_Cβ_, K_Cγ_, LC_δ2_ signals) show higher signals in the direct excitation spectrum compared to their intensities in CP spectrum (Figure 2A top).

**Figure 2.**
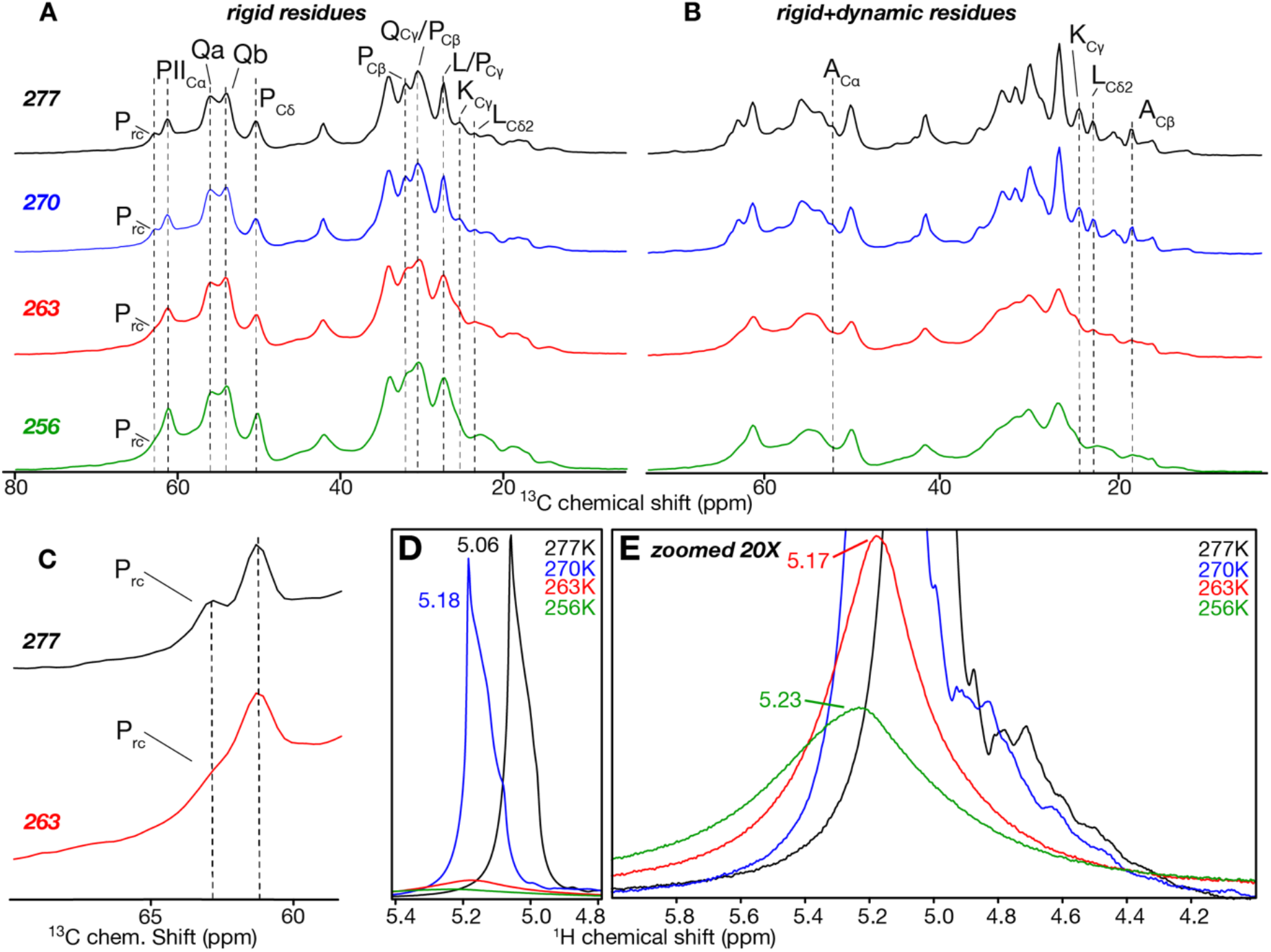
Variable temperature 1D MAS ssNMR of Q44-HttEx1 fibrils. (A) 1D ^13^C CP-MAS ssNMR at 277, 270, 263 and 256K, reflecting signals of rigid and immobilized parts of the fibrils. Dashed lines mark select peaks in different parts of HttEx1, as indicated. (B) 1D ^13^C direct excitation MAS ssNMR showing signals of both (semi)rigid and mobile residues. All data acquired at 600 MHz (^1^H) and 10kHz MAS rates. Assignments are based on (Lin et al., 2017). (C) Enlargement of Cα signals for random coil (rc) and PPII prolines. (D) ^1^H NMR of water molecules at 277K (black), 270K (blue), 263K (red), 256K (green), as indicated. (E) Same data as panel D, zoomed 20X to show low intensity signals at 263 and 256K.

Upon freezing of the surrounding water, we expected dynamics of the protein to reduce, with a pronounced impact on solvent-interacting surface-exposed residues (Mandal et al., 2015; Mandal and van der Wel, 2016). By ^1^H NMR (see below) we detect that sample water freezes between 270 and 263K (Figure 2D-E). At this temperature we observe an increase in the ^13^C CP signal for the exposed flanking segments, which normally show reduced intensities due to dynamic averaging of the dipolar interaction (Figure 2A; 256K). Signal intensity increased in the methyl group region (10-25 ppm) and for the proline signals (marked with “P”). The signals were also broadened at low temperature, reflecting a freezing out of heterogeneous conformations. Peaks from the polyQ domain also increased in intensity upon freezing (Figure S1 A). This was surprising, since the multi-nm-wide β-sheet fiber core is devoid of water and highly rigid (Schneider et al., 2011; Sivanandam et al., 2011; Hoop et al., 2016; Boatz et al., 2020). Given the stability and rigidity of the core proper, we attribute the Gln CP signal increase to an immobilization of semi-rigid glutamine residues on the fibril-core surface, as we discuss further below.

The ssNMR signals from the fiber core show equal amounts of two types of glutamine (“Qa” and “Qb”; **Figure 2**A) and reflect two differently structured β-strands in the Q44 β-hairpin (Hoop et al., 2016). Deconvoluted signals of the aliphatic glutamine carbons are shown in Figure S2 A-D, and signal widths and peak areas are indicated in Table S2. The line width of the Qa signal was unaffected by freezing, and the Qb signal showed only small changes in width. This shows that these glutamine signals reflect residues with high degree of inherent order, with little evidence of a freezing out of a broad range of disordered conformations (Table S2).

The Pro signals from the PRD underwent more pronounced changes upon sample cooling. A clear increase in the CP signal (Figure S1 A) indicates a notable change in mobility at low temperatures. As highlighted in Figure 2C, there are distinct signals from polyproline II (PII) and random coil prolines (Prc) (Lin et al., 2017). These two populations are most easily distinguished by the Cα signals (marked PII_Cα_ vs Prc) just above 60 ppm. A big cooling-induced increase in intensity was observed for the PII_Cα_ peak (Figure S1), whilst at 263K the signal of the random coil prolines, Prc, was no longer well resolved (**Figure 2**C). This finding is consistent with the prior conclusion that Prc signal reflects flexible residues (Caulkins et al., 2018; Lin et al., 2017) which can be trapped in a broad range of conformations leading to extreme signal broadening (Figure S2, Table S2). In contrast, only modest increases affected the line width of PII_Cα_ (0.2 to 0.3 kHz), showing that the PPII helices are dynamic, but also internally ordered. These findings provide additional support to previous studies showing distinct levels of order in the different parts of the PRD (Bugg et al., 2012; Caulkins et al., 2018; Falk et al., 2020; Lin et al., 2017).

### Solvent interactions and dynamics

To directly assess the connection between the changes in protein motion and solvent dynamics, we used ^1^H-detected MAS ssNMR to monitor the solvent mobility as a function of temperature (Böckmann et al., 2009; El Hariri El Nokab et al., 2022; Mandal and van der Wel, 2016). **Figure 2**D shows the 1D proton spectra of the same fibril sample between 277 and 256K. At 277 and 270K the ^1^H signal was strong, indicating rapid motion for the majority of detected water molecules. The signal shifts from 5.06 to 5.18ppm (between 277 and 270K), consistent with the temperature dependence of the water chemical shift (Schmidt et al., 2006). At 263K the signal intensity drastically lowered by ∼70% compared to the 277K signal (Figure 2D red) indicating a phase transition of bulk water from liquid to solid. A residual water signal at 5.17 ppm (Figure 2E) was from a subpopulation of water molecules that resist freezing, but did have a broadened peak width demonstrating constrained mobility. These are expected to be water molecules close to the protein (Böckmann et al., 2009). At 256K the water signal weakened by ∼75% compared to the original intensity at 277K and broadened further, indicating a progressive immobilization of the water molecules close to fibrils. Overall, these ^1^H NMR results demonstrate the different dynamics of different pools of solvent molecules around the HttEx1 fibrils. The coincident behavior of the ^1^H and ^13^C NMR changes support the idea that the changes in motion of the protein core interface and flanking domains (seen by ^13^C NMR) are coupled to the water motional changes.

### 2D ssNMR of local fibril order and disorder

A more detailed analysis of the temperature-related changes in the protein was obtained from 2D ssNMR experiments at 277 and 256K, above and below the solvent freezing temperatures (**Figure 3**A, B). The two spectra have many features in common. The dominant ssNMR signals of a- and b-type glutamines were easily recognized at both temperatures (marked red (‘a’-type Gln) and blue (‘b’ type Gln)). The line widths of these polyQ-core signals Qa and Qb showed only negligible changes between high and low temperature (**Table 2**). This finding is consistent with their participation in the fiber core, with limited contact with the (freezing) solvent. At 277K a much weaker signal from a third type of glutamines (Qc in Figure 3A, grey box) was observable, previously attributed to non-core glutamines (Hoop et al., 2016; Isas et al., 2017, 2015; Sivanandam et al., 2011). At 256K, this signal broadened and disappeared, supporting the idea that it must stem from residues interacting with the solvent, pointing to a presence outside the internal fibril core (Helmus et al., 2008).

**Figure 3.**
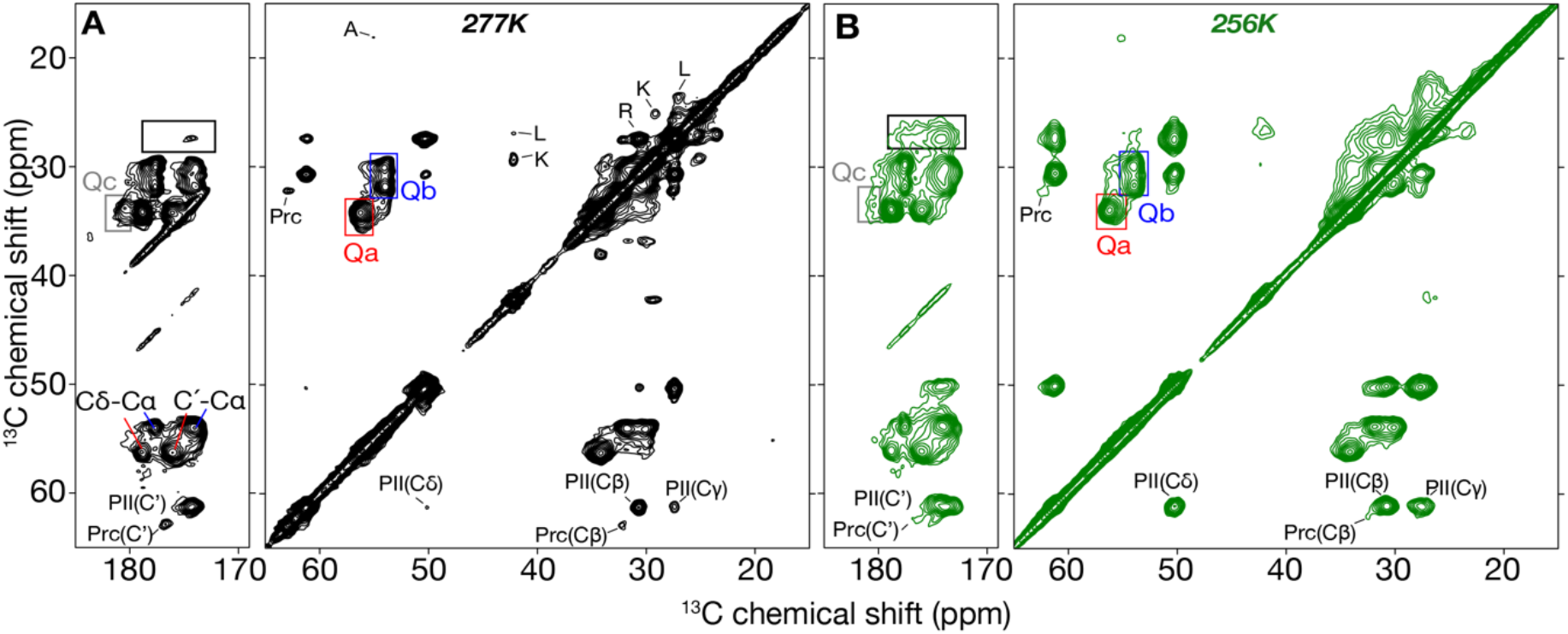
2D ^13^C-^13^C DARR of Q44-HttEx1 fibrils. (A) 277K and (B) 256K. Cross-peaks for the backbones of the core glutamine types “a” (Qa) and “b” (Qb) are marked with red and blue boxes respectively. Their C’-Cα and Cδ-Cα peaks are labeled in panel A. The Cδ-Cγ peak for a third (non-core) glutamine type, Qc, is shown in grey box in panels A and B. A black box marks CO-Cδ peaks for proline residues. The spectra were acquired with a 25 ms DARR mix time at 10 kHz MAS.

**Table 2.** Linewidths extracted from 2D experiment for two ‘a’ and ‘b’ glutamine types. Linewidths of 2D peaks from the CP/DARR ^13^C-^13^C spectra measured at 277 and 256K.

| Residues | Qa line widths (kHz) |  | Qb line widths (kHz) |  |
| --- | --- | --- | --- | --- |
|  | 277K | 256K | 277K | 256K |
| Cα | 0.12 | 0.10 | 0.12 | 0.10 |
| Cβ | 0.17 | 0.18 | 0.18 | 0.19 |
| Cγ | - | - | 0.18 | 0.19 |
| Cδ | 0.16 | 0.18 | 0.13 | 0.14 |
| C’ | 0.18 | 0.23 | 0.18 | 0.19 |

Unlike the polyQ core, the PRD segment’s proline signals display a notable peak broadening in the 2D spectra, with different effects on the PPII helix and random coil prolines (Figure 3A, B, PII, Prc). The PPII cross peaks become stronger with temperature reduction, while the cross-peaks of Pro in the random coil conformation became weaker. The effect is even better seen in long range magnetization transfer at 256K where Prc signals totally disappeared, but PPII became stronger (Figure S4). Additional backbone and side chain correlations were observed between prolines in the PPII helix conformation (similar to the polyQ core peaks). Thus, the PPII and Prc prolines experience distinct mobilities. Referring also to the 1D data analysis (above), we propose that the random coil signals experience a dramatic broadening. Moreover, given that part of the fiber-bound water (see above) retains partial mobility, the C-terminal tail may also retain some continued dynamics even at the lowest measurement temperature.

From prior work we also know the spectral locations of peaks from the (partly α-helical) Htt^NT^ segment (Lin et al., 2017). Such signals were observed in the 1D and 2D spectra at 277K, and include well-resolved peaks from various methyl-bearing aliphatic amino acids (Ala, Met, Leu), as well as Lys (typical ^13^C shifts between 10 and 35 ppm. Upon sample freezing, many such cross-peaks near the 2D diagonal between 38 to 22 ppm became less apparent due to excessive broadening (Figure 3B). These observations contrasted with the signals of the more ordered polyQ and even PPII proline residues, and reinforce the idea that the Htt^NT^ segment may have a partial secondary (α-helical) structure but also a high degree of dynamic disorder.

### Variable temperature 2D ^15^N-^13^C spectra

2D double cross-polarization (DCP) experiments based on ^1^H-^15^N and ^15^N-^13^C CP steps were performed at 277 and 256K. At ambient temperatures, increased backbone mobility impacts both CP steps leading to losses of signal intensity for dynamic residues, as described e.g. for aggregated prion proteins (Helmus et al., 2008). Figure 4A,B compares the 2D NCO and NCA spectra of Q44-HttEx1 fibrils at 277 and 256K. At 277K, the spectra are dominated by signals from the polyQ amyloid core, with no signals from the flanking segments. The lack of signal for the latter stems from a combination of solvent-coupled dynamics and the lack of N-H protons in the prolines. The impact of low temperatures on these spectra is more striking than in the ^13^C-^13^C 2D spectra. At low temperature many additional signals were detected in NCA spectrum, showing the improved polarization transfer in absence of molecular motion (Figure 4B). Many of these signals are very broad, including an observed glycine signal (only present in the PRD; Table 1). For the polyQ core, we also now observe longer range transfers (e.g. N-Cγ signals), revealing a lack of specificity in the configuration of the SPECIFIC CP used in the NCA experiment. Although this could impact detailed quantitative features in the spectra, we here focus on a qualitative comparison of relative linewidths of observed peaks. Also the PPII prolines become visible due to improved long-range polarization transfer overcoming their lack of protonation of the backbone nitrogen. Both sidechain and backbone PPII signals had widths of 0.17 and 0.19 kHz, indicating a well-ordered PPII helical structure (Figure 4B).

**Figure 4.**
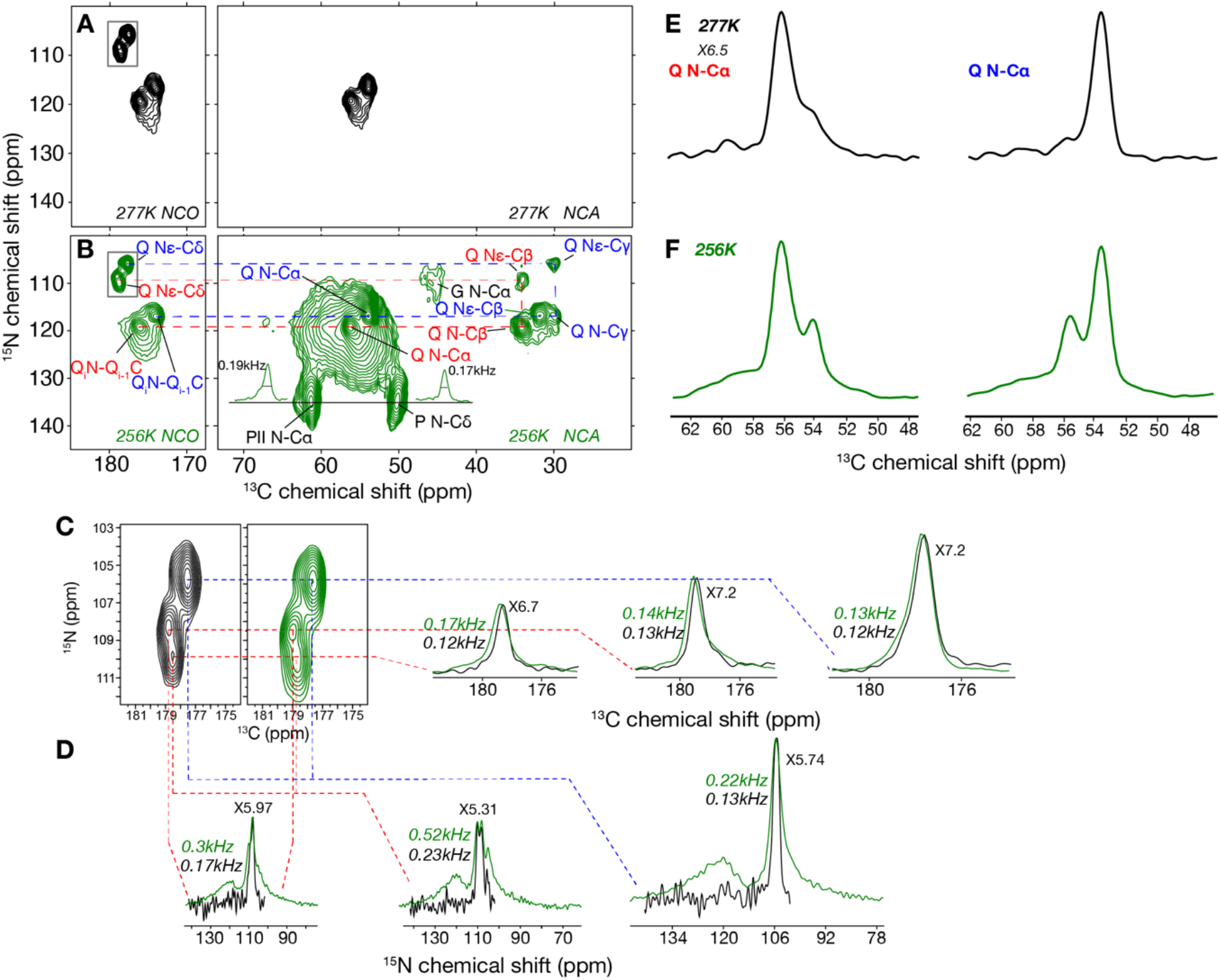
2D N-C spectra at 277 and 256K. (A) NCO and NCA spectra at 277K and (B) at 256K. (C) Zoomed Q Nε-Cδ connectivities shown in box in panels A and B at 256K (green) and 277K (black). Their 1D horizontal slices show small changes in ^13^C signal widths. (D) 1D vertical slices of panel (C) show larger discrepancies in ^15^N signal line widths between two temperatures. (E) 1D ^13^C N-Cα slices of type a and b Gln at 277K extracted at ^15^N = 119.3 ppm and 116.9 ppm. (F) 1D N-Cα slices at 256K for ^15^N = 119.3 and 116.9 ppm. The signal intensity in E was increased 6.5 times to reach same intensity as at 256K. In all panels, red labels and lines indicate type “a” Gln and blue labels and lines refer to type “b” Gln.

At low temperature, the 2D N-C spectra gained a broad background signal surrounding the consistently narrow peaks of the polyQ core residues (Figure 4B-C). Figure 4 also shows selected 1D slices through these 2D spectra, to allow a clearer view of the differences in linewidth. We see small increases in line widths of the side chains of both glutamine types, for both the ^13^C and ^15^N dimensions (Figure 4C,D). The highest increase was observed for type “a” glutamine (from 0.12 to 0.17 kHz for ^13^C). These changes indicate that the glutamine side chains experienced the solvent freezing. The 1D slices of backbone signals (N-Cα) are shown in Figure 4E,F. At both temperatures, signals were deconvoluted using three peaks. Both Qa and Qb maintain their linewidths at two temperatures indicating that these glutamine backbones are highly ordered in the fiber core and relatively insensitive to the solvent mobility. However, a broad third signal at 59.5ppm (Figure 4E) accounts for 7.75% of total integrated Cα signal intensities. At 256K, this signal becomes ∼16% of the integrated intensity and shifts upfield to 58.3ppm (Figure 4F). This region of the spectrum is expected to show signals from most HttEx1 residues except the prolines. Our Q44-HttEx1 construct contains in total 120 residues composed of 31 prolines, 50 glutamine residues, and 39 other residue types (**Table 1**). Thus, up to 39 non-polyQ residues could contribute to the broad signals, along with a limited number of polyQ residues that are outside the core and free to adopt a broad range of conformations. However, while visible in these low-temperature spectra, the peak broadness means that they are impossible to separately study or even identify.

### Temperature-dependent changes in dipolar order parameters

We observed an increase in (CP ssNMR) signal intensity for different residue types, coinciding with solvent freezing. This points to these residues normally experiencing solvent-coupled motion, suggesting a location outside the internal fibril core. Partly these were highly flexible residues that freeze into very broad signals in a frozen state. Other amino acids retain relatively narrow signals, which increase in intensity upon freezing, but show limited broadening. We attribute these to residues (especially glutamines) that reside on the outer surface (“core-water interface”) of the polyQ fiber core. These residues would disply partial mobility, such that they appear in CP spectra but have reduced dipolar couplings and thus diminished signal intensity. Such peaks are difficult to observe, especially if they are close to (or overlap with) dominant peaks of the rigid polyQ core. We therefore explore an approach for selectively probing such semi-rigid residues, with CP-based ssNMR experiments in which rigid sites are selectively eliminated based on their strong ^1^H-^13^C dipolar couplings.

The impact of molecular motions that modulate dipolar interactions can be measured via dipolar chemical shift correlation (DIPSHIFT) CP-ssNMR experiments (Munowitz et al., 1981). Figure 5A illustrates this for simulated ^13^C-^1^H dephasing curves where rapid dephasing represents more order (less motion) and slow dephasing indicates increased mobility. The corresponding experimental dephasing curves were acquired for unfrozen and frozen Q44-HttEx1 fibrils (**Figure 5**B-C). Both types of glutamines in the fiber core showed rapid dephasing (and clear dipolar oscillations) at 277K, indicating a similarly high backbone rigidity even without sample freezing (**Figure 5**B). The obtained data were consistent with prior measurements performed under different conditions or using different fibril types (Hoop et al., 2014; Lin et al., 2017). On the other hand, proline signals show slow dephasing (**Figure 5**D) indicating a higher mobility (low order parameters). In addition, prolines with random coil structure experienced higher mobility than those in the PPII helices. Note that all these measurements rely on CP-based ssNMR, which means that the detected residues lack rapid isotropic motion.

**Figure 5.**
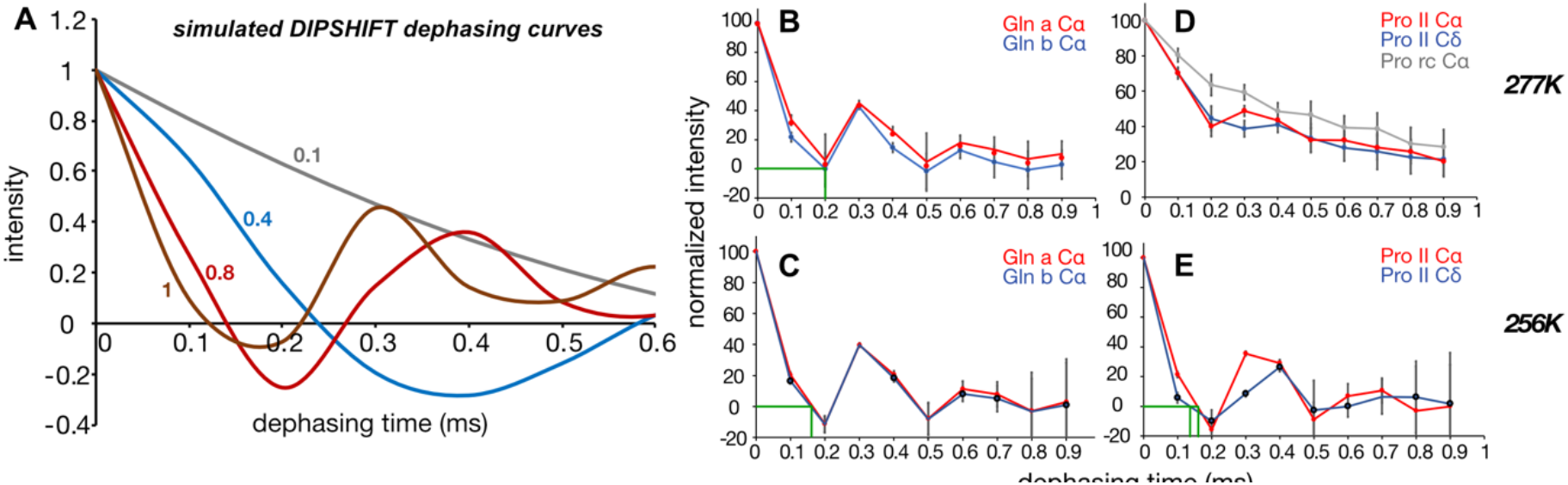
Dipolar dephasing curves for Q44-HttEx1 at variable temperature. (A) Simulated ^13^Cα-^1^H dephasing curves based on R18 ^7^ -based ^1^H-^13^C dipolar recoupling, for different order parameter values between 0.1 and 1 and with 0.5ms relaxation (Lin et al., 2017). (B) Experimental dephasing curves of the ‘a’ and ‘b’ glutamine types at 277K, and (C) at 256K. (D) Experimental dephasing curves of prolines from PPII helices (red, blue) and from the random coil conformation (grey) at 277K, and (E) at 256K. Experimental data were measured at 600 MHz (^1^H) and 8 kHz MAS.

Upon freezing the sample (at 256 K), the Pro dephasing curves became similar to those of the rigid amyloid core. Notably, also the polyQ signal dephasing became more pronounced than at 277 K, indicating a higher dipolar coupling constant. The polyQ dephasing curves cross the x-axis at ∼0.17 ms recoupling time, whilst Pro Cδ and ProII Cα reached zero signal at 0.14 and 0.16ms, respectively (**Figure 5**E). Note that the polyQ dephasing reached zero at 0.20m ms at 277K. Thus, again we observe a close connection between solvent freezing and the segment-specific dynamics of the HttEx1 fibrils, with signs of a solvent-coupled change also affecting the β-sheet polyQ segment signals.

### Revealing residues with intermediate dynamics

Next, we used DIPSHIFT dephasing to selectively suppress the normally dominant signals from the rigid and dehydrated fiber core, which can obscure weak signals from more mobile (surface) residues. As noted above, in the unfrozen sample a dephasing time near 0.2ms caused the most rigid residues to dephase (signal intensity zero or negative) while more mobile residues were still observable with positive signal intensity (Figure 5). Thus, by combining CP-based signal generation with ^1^H-^13^C DIPSHIFT recoupling one can obtain spectra showing primarily those residues with intermediate dynamics, yielding an intermediate motion selective (IMS) DYSE experiment. We combined a DIPSHIFT dephasing block with a canonical ^13^C-^13^C DARR pulse sequence (**Figure 6**) to test this approach on the same Q44-HttEx1 fibril sample.

**Figure 6.**
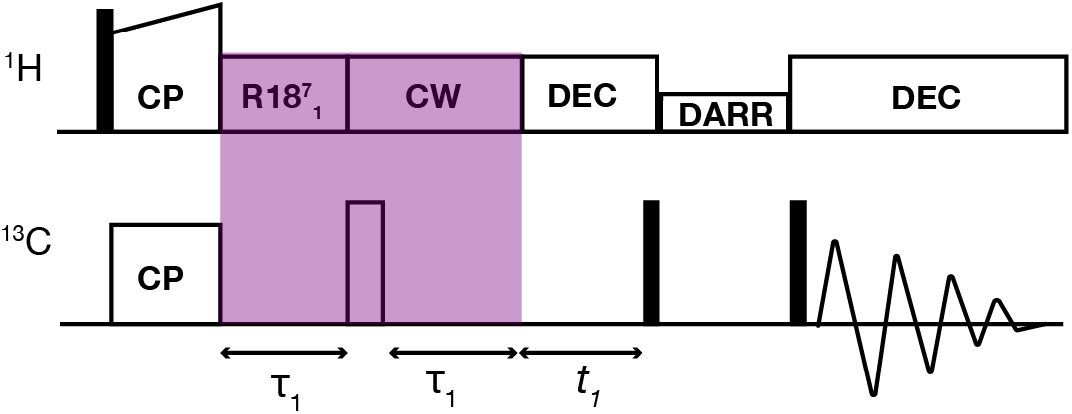
Pulse sequence of combined DIPSHIFT and DARR experiments. After the ramped CP, a DIPSHIFT dipolar filter is applied, with a fixed ^1^H-^13^C recoupling period, here shown with R18_1_^7^ recoupling. Subsequently, the residual ^13^C signal (from non-rigid carbons) is allowed to evolve under incrementing t_1_ periods, prior to DARR-based ^13^C-^13^C polarization exchange. This pulse sequence generates a broad-band ^13^C-^13^C spectrum, but the DIPSHIFT filter would only work fully for the ^13^C-^1^H protonated carbons (CH/CH_2_).

A normal ^13^C-^13^C DARR spectrum without dynamic filtering is shown in **Figure 7**A, whilst the dynamically filtered variant is shown in **Figure 7**B. After 0.2ms of DIPSHIFT filtering, the most rigid residues appear as negative signals (orange) whereas positive cross-peaks (blue) represent residues that have not been dephased completely. Therefore, these latter signals stem from partly mobile sites, e.g. residues outside the buried and dehydrated fiber core itself (ref. Figure 1C). Note that the normal 2D spectrum is dominated by signals from the rigid polyQ core residues, and that these were indeed effectively eliminated by the tuned DIPSHIFT filter. Here, the DIPSHIFT experiment was optimized for dephasing of protonated, immobile aliphatic carbons (in particular C-H groups). Methyl groups and non-protonated carbons (e.g. C=O groups) were not expected to be properly dephased, even in cases where they are part of the rigid structure. Off-diagonal peaks involving the polyQ core (type ‘a’ and ‘b’) Cα appear as negative peaks, indicating their rigidity. This is true for Cα-to-C=O cross-peaks as well. The filtering is applied prior to the t_1_ evolution, such that the “filtered” C-H group should be along the indirect dimension of the 2D spectrum. The converse signal (from C=O to Cα; across the diagonal) is not negative in intensity (Figure S4).

**Figure 7.**
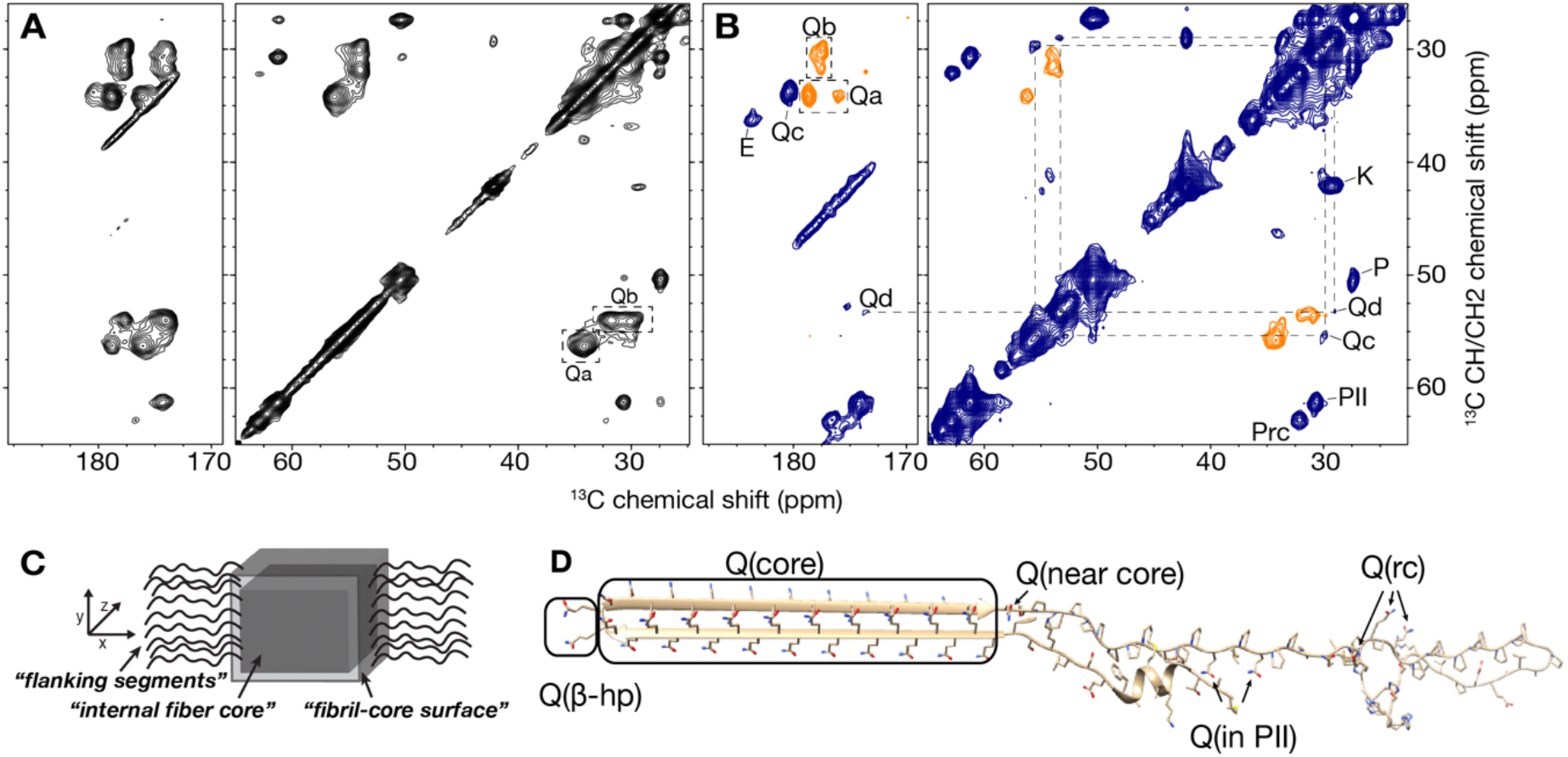
Intermediate-motion selection DYSE in ^13^C-^13^C 2D MAS NMR. (A) 2D ^13^C-^13^C DARR experiment at 277K. The dominant Cα-Cβ/Cγ peaks of the rigid polyQ core signals “Qa” and “Qb” are marked with dahs boxes (right). (B) Corresponding DIPSHIFT-filtered DARR experiment at 277K, using a 0.20 ms R18_1_^7^ dipolar recoupling period at 10 kHz MAS. The previously marked rigid core signals now are negative (orange), while previously invisible glutamine signals become visible as positive peaks (blue). These are marked as Qc and Qd. Other strong positive signals from non-Gln residues outside the fiber core are also marked (E, K, P residues). The amino acid type assignment and secondary structure analysis was done using PLUQ program (Fritzsching et al., 2016, 2013). (C) Schematic drawing of the fibril structure, showing the nomenclature distinguishing the internal fiber core (i.e. dry interfaces) and segments and residues outside the core (wet interface; flanking domains). (D) Schematic monomer in our model of the Q44-HttEx1 fibril structure, marking the locations of glutamine residues outside the dry fiber core: β-turn residues labelled as Q(β-hp), terminal polyQ segment residues (“near core”), and a few Gln in the dynamic PRD indicated as “rc” and “PII”.

The purpose of this experiment was to identify and observe semi-mobile residues. These peaks would manifest as positive off-diagonal peaks (shown in blue) in the chemical shift range of non-methyl aliphatic carbons (70 to 30 ppm on the vertical axis). Indeed, we note several such peaks, with the strongest signals stemming from the proline residues of the PRD (marked Prc and PII). These signals had been weak to modest in the normal DARR spectrum (**Figure 7**A), but were prominent in the filtered spectrum. Notably, the ‘random coil’ Pro signal (Prc) was particularly strong relative to the PII peak, compared to the normal DARR 2D. This illustrates nicely that the signal intensities are dependent on the motional properties. It is important to note that this implies that the observed signal intensities should not be seen as quantitative indicators of the relative populations of the detected residues. Rather the purpose is to detect otherwise invisible species, normally masked by more rigid signals.

We had a particular interest to probe glutamine residues at the core-water interface, which have chemical shifts similar to those of the rigid internal fiber core. Partial insight into these residues was reported in prior work, but a detailed study was complicated by the proximity of dominant rigid core signals. In the filtered spectrum, previously invisible positive glutamine signals now became observable, close to the (orange) filtered signals of the internal fiber core. The signal at 29.9 × 55.5 ppm lines up well with Qc form (**Figure 7**B), previously identified as more dynamic Gln residues (from site-specific labeling studies). These chemical shifts of Qc in the full exon 1 resemble the Q47c1 signal site-specifically assigned in residue-specifically labeled htt^NT^Q_30_P_10_K_2_ Htt N-terminal fragment peptide where it reflected a non-β-strand configuration just outside the amyloid core (Hoop et al., 2014) (**Figure 7**C, Q(near core); Table S3). A second positive signal appears at 29.0 × 53.3 ppm. Correlation of this signal to carbonyl region (Cα-to-C=O) at 173.6×53.3ppm appears as a weak positive signal as well. The PLUQ program (Fritzsching et al., 2016, 2013) for amino acid assignment and secondary structure analysis suggests the highest probability (∼90) of this signal stemming from Gln residues with a random coil conformation, or a low probability (∼10) for His residues. We deem the latter unlikely, given that the only His residues are in the C-terminal His-tag (Table 1), and that this His tag (in our fibrils) is flexible and not detected in CP-based spectra. Thus, we attribute the cross-peaks at 29.0 × 53.3 and 173.6 × 53.3 ppm to a distinct glutamine population (coined Qd) with a random coil conformation (**Figure 7**B). A small number of Gln residues is also present in the flanking segments, specifically, in the C-terminal PRD tail (Figure 7C), which is typically more visible in INEPT-than in CP-based spectra (Caulkins et al., 2018; Lin et al., 2017). As these DIPSHIFT-DARR measurements depend on CP ssNMR, we attribute these semi-rigid Gln signals to residues that are part of the polyQ segment, but not buried: solvent exposed yet semi-rigid due to being proximal to the rigid polyQ core.

## Discussion

The current work presents complementary approaches to dissect the complex dynamics of Q44-HttEx1 fibrils, which feature dynamics spanning from highly rigid to highly flexible. This range of molecular motion is common to many pathogenic protein fibrils, including tau and α-synuclein (Andronesi et al., 2008; Fitzpatrick et al., 2017; Helmus et al., 2010, 2008; Tuttle et al., 2016). Dynamically disordered flanking segments are biologically interesting as they appear to be target sites for disaggregating chaperones and impact fiber toxicity and polymorphism (Chongtham et al., 2020; Duennwald et al., 2006; Lin et al., 2017; Robertson et al., 2010; Wentink et al., 2020). However, good experimental methods to study all parts of these dynamic regions have been missing.

Our variable-temperature measurements on the protein fibrils showed how changes in the solvent dynamics impacted the mobility of different regions of the HttEx1 fibrils. Solvent freezing leads to an extreme broadening of signals from the most flexible flanking domain residues, which in some spectra effectively prohibited visibility. Highly broadened signals likely stem from the htt^NT^ and PRD flanking segments, which are almost entirely devoid of β-sheet structure and are solvent exposed and dynamic (Sivanandam et al., 2011). However, the noted signal broadening renders a detailed analysis difficult to impossible, even if the signals are clearly visible in the low-temperature spectra. Thus, we see large limitations to the use of low-temperature experiments to study the fibrils’ flexible flanking segments and other solvent-exposed residues. That said, one intriguing observation in the low temperature data was that the impact on the Gln backbone and side chain resonances is very small. This can only be explained by these structural elements being remote from the solvent phase, and insensitive to its mobility. This observation is well explained by our previously published structural model of the polyQ core of these HttEx1 fibrils, reproduced in Figure 1A (Boatz et al., 2020). It shows a fiber core containing approximately nine interdigitating β-sheets, placing the Gln side chains in a constrained steric-zipper arrangement. In this core structure, a large majority of the Gln residues is placed in the internal fiber core, such that one may estimate a maximum of 19 % of the residues in the polyQ segment exposed to solvent (i.e. in turns/loops and in solvent-facing β-strands such as those shown yellow in Figure 1A). Interestingly, upon sample freezing, the signals of the polyQ core do display a modest change in certain parameters, such as the dipolar order parameter and linewidth. This effect is most notable for the Gln side chain signals. We propose that this may stem from a population of solvent-exposed polyQ residues that compose the core-water interface of the layered β-sheet core (Figure 1). There may also be a contribution from a freezing of residual motion throughout the polyQ core, but we suggest that this is a less critical factor given that the core is assembled from multiple layers of β-sheets (based on ssNMR and EM studies; (Boatz et al., 2020; Nazarov et al., 2022)). This renders the dehydrated core multiple nanometers thick, creating a more extensive and rigid core structure than present in most amyloid proteins.

The glutamine residues facing the solvent on the fiber surface are of significant interest. They include the as-yet poorly understood β-turn regions of the β-hairpins of polyQ-expanded HttEx1 fibrils (Hoop et al., 2016; Matlahov and van der Wel, 2019). Furthermore, the fibril-core surface has surface grooves that are characteristic of amyloid-like fibrils, and feature the binding sites of both amyloid-specific fluorescent dyes and other diagnostic tracer molecules (Bertoglio et al., 2021; Biancalana and Koide, 2010; Fändrich et al., 2018; Schütz et al., 2011). Yet, in normal ssNMR spectra these solvent-facing glutamine residues are hard to distinguish from those internal to the polyQ core, with the latter dominating the CP-based 1D and 2D ssNMR spectra. We observed that the CP-signals from the polyQ core increase upon freezing, while retaining much of the narrowness (and chemical shift). This hints that we observe a gain in signal from residues that are solvent-mobilized but occupy a minimum in the folding landscape that closely resembles the conformation of the core proper. The observed changes in signal intensity, linewidth and dipolar dephasing at low temperature provide information on solvent-interacting segments and residues, but the level of detail is limited. We previously reported some similar analyses in related but different polyQ peptide fibrils: we analyzed the motions of site-specifically labeled residues in the polyQ and Htt^NT^ segments of Htt N-terminal fragment peptide fibrils (Hoop et al., 2014). However the current study is the first report of such measurements applied to HttEx1 fibrils.

As a complementary new approach we used dipolar dephasing to introduce (to the best of our knowledge) a new DYSE ssNMR measurement, which is focused on observing signals from residues with a mobility in-between highly rigid and highly flexible. A similar technique was recently reported using a different approach via ^13^C-^15^N REDOR filtering to selectively observe protein signals displaying ms time scale motion (Kashefi et al., 2019). In our approach, we integrated a DIPSHIFT-based filtering element in a standard CP-ssNMR pulse sequence, to selectively dephase signals from rigid protein segments, thus highlighting the signals from semi-rigid residues. We show how this approach allows us to observe previously invisible cross-peaks from semi-rigid residues, which in normal spectra are overwhelmed by nearby high-intensity signals from the internal fiber core structure. The observed resonances reveal two semi-rigid types of peaks, with chemical shifts indicative of Gln residues (summarized in Table S3). One of these peaks has resonances previously detected for site-specifically labeled Gln residues (Hoop et al., 2014, 2016; Kar et al., 2017). The other (type “d”) signals are distinct and are tentatively attributed to another type of semi-rigid glutamine. Though distinct from each other, both have chemical shifts that are inconsistent with Gln in β-sheet structure, as expected for non-amyloid-core residues. Notably, detecting solvent-facing Gln residues is of interest as they may form surface features of mutant HttEx1 fibrils that are recognized by fluorescent dyes, PET ligands, and certain types of HttEx1 aggregation inhibitors (Bertoglio et al., 2021; Kar et al., 2017; Matsuoka et al., 2012; Minakawa and Nagai, 2021). A further characterization of the structural features of these residues requires additional studies, which are ongoing.

The described IMS DYSE experiment has a number of limitations or alternative applications. First, in the current example we focused on filtering CH and CH_2_ groups, noting that other sites (e.g. CO, CH_3_) would not be optimally affected. It may be feasible to tune the DIPSHIFT filter specifically for CO or CH3 groups, but in that case the experiment would not be effective for the CH/CH_2_ groups. So it is in a sense a narrow-banded experiment. As demonstrated here, we obtained negative peaks for rigid signals at the chosen dephasing time. Depending on the experimental goals, it may be preferable to tune the MAS rate and mixing time to achieve perfect elimination (i.e. exactly zero, instead of negative signal) of rigid sites, which may enable detecting of strongly overlapping peaks.

DIPSHIFT-filtration of C-H signals also drastically decreases overall spectrum sensitivity. As a result, weak signals with desired mobility may disappear from the spectrum. Secondly, it is impossible to perform quantitative analysis of the detected populations of (mobile) residues, because the peak intensity scales not only with the number of residues (or atoms) detected, but is also directly dependent on the dynamic features. Secondly, a single DYSE spectrum does not allow the dipolar order parameters to be determined. These limitations may narrow the range of systems or conditions where this experiment can be most useful. However, akin to other DYSE measurements, we anticipate this approach to be a tool for assessment of different dynamics in systems with resolved and (partly) overlapping peaks. This measurement can be complementary to more established techniques to probe highly flexible residues. Such protein regions are routinely studied using INEPT-based ssNMR approaches. For instance, prior studies have reported extensively on the flexible parts of the C-terminal tail of HttEx1 based on such experiments (Boatz et al., 2020; Caulkins et al., 2018; Isas et al., 2017, 2015; Lin et al., 2017).

The use of the dipolar-filter to selectively suppress signals from rigid components and observe semi-rigid sample components, could also be extended to other types of ssNMR experiments. The described integration with a broadband technique such as the employed DARR ^13^C-^13^C 2D spectrum works, but one needs to keep in mind that it designed specifically for non-methyl protonated aliphatic carbons (CH, CH_2_ groups). Therefore, an attractive use may be to integrate it into Cα-selective experiments, such as NCA(CX) pulse sequences. However, as discussed around Figure 4, a caveat with such an approach is that such DCP experiments suffer from a limited recovery of signals from (semi) mobile residues. A workaround for this issue may be the use of a (frequency selective) TEDOR sequence to measure the equivalent of an NCA experiment (Daviso et al., 2013) with intermediate mobility selection. Naturally, many other variations in implementation could be designed, for instance toward applications at fast MAS, alternative ^1^H-^13^C recoupling techniques, etc.

We envision useful applications of IMS-DYSE experiments beyond amyloid fibrils. In many functional proteins, loops, turns and even whole (enzymatic) domains undergo millisecond timescale motion that may be essential for protein function. Such structural re-arrangements of active sites depend on interactions with water molecules, with hydration and motion being essential for protein function (Singh et al., 2019; Vasa et al., 2019; Klein et al., 2022). This applies to many proteins studied by ssNMR, including crystalline globular proteins, large protein complexes, and membrane proteins (Eddy et al., 2015; Kashefi et al., 2019; Saurel et al., 2017). Thus, hard-to-detect NMR signals from enzymatic active sites and membrane proteins can be a target for such DIPSHIFT-based filtering techniques.

## Conclusions

We employed complementary methods in ssNMR to study the dynamics of Q44-HttEx1 fibrils, focusing on segments or residues outside but near the rigid fiber core. Such residues have previously remained invisible or were only tangentially observed. The mobility was shown to be strongly correlated to the dynamics of the fiber-associated water molecules. Variable temperature studies reveal the presence of dynamic components, but fail to give detailed insights due to the excessive signal overlap. As a work-around we described and applied a new ^1^H-^13^C DIPSHIFT-based dynamic-based spectral editing experiment that suppresses signals from the rigid core, and emphasizes the CP-visible signals of partly dynamic core-proximal protein segments. This technique revealed two normally masked cross peaks of semi-rigid residues, by suppressing a large excess of rigid polyQ core signals. These methods and results pave the way for further investigations of the fiber surface, with potential implications for diagnostic and inhibitor developments relevant to HD. Moreover, we anticipate similar methods to be useful to a variety of other protein fibrils, membrane proteins, and (bio)materials with solvent-facing chemical groups attached to a rigid core. For instance, IMS-DYSE measurements could enhance ssNMR studies of nanoparticles, nanomaterials, thin films and bottlebrush polymers (Golkaram et al., 2019; Gruschwitz et al., 2020; Protesescu et al., 2014; Walsh and Knecht, 2017).

## Acknowledgements

This work was supported by the National Institutes of Health (grants R01 GM112678 and R01 GM113908 to PvdW; T32 GM088119 to J.C.B), the Achievement Rewards for College Scientists (ARCS) Foundation (J.C.B.), the CHDI Foundation, and the CampagneTeam Huntington foundation.

## Abbreviations

AD: Alzheimer’s disease
CP: cross polarization
cryo EM: cryogenic electron microscopy
DARR: dipolar assisted rotational resonance
DCP: double CP
DIPSHIFT: dipolar chemical shift
DYSE: dynamic spectral editing
HD: Huntington’s disease
HttEx1: huntingtin exon 1
htt^NT^: N-terminal domain of htt
IMS-DYSE: intermediate-motion selection dynamic spectral editing
INEPT: insensitive nuclei enhanced by polarization transfer
MAS: magic angle spinning
MBP: maltose-binding protein
MBP-Q44-HttEx1: maltose-binding protein bound to huntingtin exon 1 which includes 44 glutamines in its polyglutamine domain
PD: Parkinson’s disease
PDSD: proton-driven spin diffusion
PET: positron emission tomography
polyQ: polyglutamine
PPII: polyproline II
PRD: proline-rich domain
Prc: prolines in random coil conformation
REDOR: rotational echo double resonance
ssNMR: solid-state nuclear magnetic resonance
T_2_: transverse relaxation
TEDOR: transferred echo double resonance
TPPM: two-pulse phase modulated proton decoupling

