## Supporting Information file for "Selective observation of semi-rigid non-core residues in dynamically complex mutant huntingtin protein fibrils"

**Table S1. Experimental parameters of MAS NMR experiments.** Abbreviations: NS, number of scans (per $t_1$ point for 2Ds); Temp., temperature; MAS, magic angle spinning rate; RD, recycle delay; Mixing, $^{13}C$-$^{13}C$ or $^{1}H$-$^{1}H$ mixing time (ms); $t_1$ evol., maximum $t_1$ evolution time expressed in number of $t_1$ points (real+imaginary) x $t_1$ increment time; CP time, cross polarization contact time (in ms).

| Experiment | Figure | NS | MAS | RD | $t_1$ evol. | Mixing | CP time | Temp |
|---|---|---|---|---|---|---|---|---|
| | | | kHz | s | ms | ms | ms | K |
| 1D $^{13}C$ CP | 2A,2C,S1A, S2A, | 256 | 10 | 3 | NA | NA | 1 | 277 |
| 1D $^{13}C$ CP | 2A,S1A, S2B | 256 | 10 | 3 | NA | NA | 1 | 270 |
| 1D $^{13}C$ CP | 2A,2C,S1A, S2C | 256 | 10 | 3 | NA | NA | 1 | 263 |
| 1D $^{13}C$ CP | 2A,S1A, S2D | 256 | 10 | 3 | NA | NA | 1 | 256 |
| 1D $^{13}C$ DE | 2B,S1B | 256 | 10 | 3 | NA | NA | NA | 277 |
| 1D $^{13}C$ DE | 2B,S1B | 256 | 10 | 3 | NA | NA | NA | 270 |
| 1D $^{13}C$ DE | 2B,S1B | 256 | 10 | 3 | NA | NA | NA | 263 |
| 1D $^{13}C$ DE | 2B,S1B | 256 | 10 | 3 | NA | NA | NA | 256 |
| 1D $^{13}C$ INEPT | S1C | 256 | 8.33 | 3 | NA | NA | NA | 277 |
| 1D $^{13}C$ INEPT | S1C | 256 | 8.33 | 3 | NA | NA | NA | 270 |
| 1D $^{13}C$ INEPT | S1C | 256 | 8.33 | 3 | NA | NA | NA | 263 |
| 1D $^{13}C$ INEPT | S1C | 256 | 8.33 | 3 | NA | NA | NA | 256 |
| 1D $^{1}H$ | 2D,2E, S3A | 8 | 10 | 2.8 | NA | NA | NA | 277 |
| 1D $^{1}H$ | 2D,2E, S3B | 8 | 10 | 2.8 | NA | NA | NA | 270 |
| 1D $^{1}H$ | 2D,2E, S3C | 8 | 10 | 2.8 | NA | NA | NA | 263 |
| 1D $^{1}H$ | 2D,2E, S3D | 8 | 10 | 2.8 | NA | NA | | 256 |
| 2D DARR | 3A,S5A,7A | 80 | 10 | 2.8 | 9.9 (624x32μs) | 25 | 1 | 277 |
| 2D DARR | 3B | 32 | 10 | 2.8 | 5.6 (414x27.025μs) | 25 | 0.75 | 256 |
| 2D PDSD | S4 | 32 | 10 | 2.8 | 5.6 (414x27.025μs) | 500 | 0.75 | 256 |
| 2D DIPSHIFT-DARR | 7B,S5B | 32 | 10 | 2.8 | 8.9 (660x27.025μs) | 25 | 0.75 | 277 |
| 1D DIPSHIFT | 5B,5D | 128 | 10 | 3 | NA | vary | 1 | 277 |
| 1D DIPSHIFT | 5C,5E | 128 | 10 | 3 | NA | vary | 1 | 256 |
| 2D NCO | 4A,4C | 128 | 13 | 2.8 | 15 (74x410) | NA | 1.5, 4 | 277 |
| 2D NCO | 4B,4C | 192 | 10 | 2.8 | 11.9 (174x137) | NA | 0.8, 4 | 256 |
| 2D NCA | 4A | 128 | 13 | 2.8 | 15 (74x410) | NA | 1.5, 4 | 277 |
| 2D NCA | 4B | 192 | 10 | 2.8 | 11.9 (174x137) | NA | 0.8, 4 | 256 |

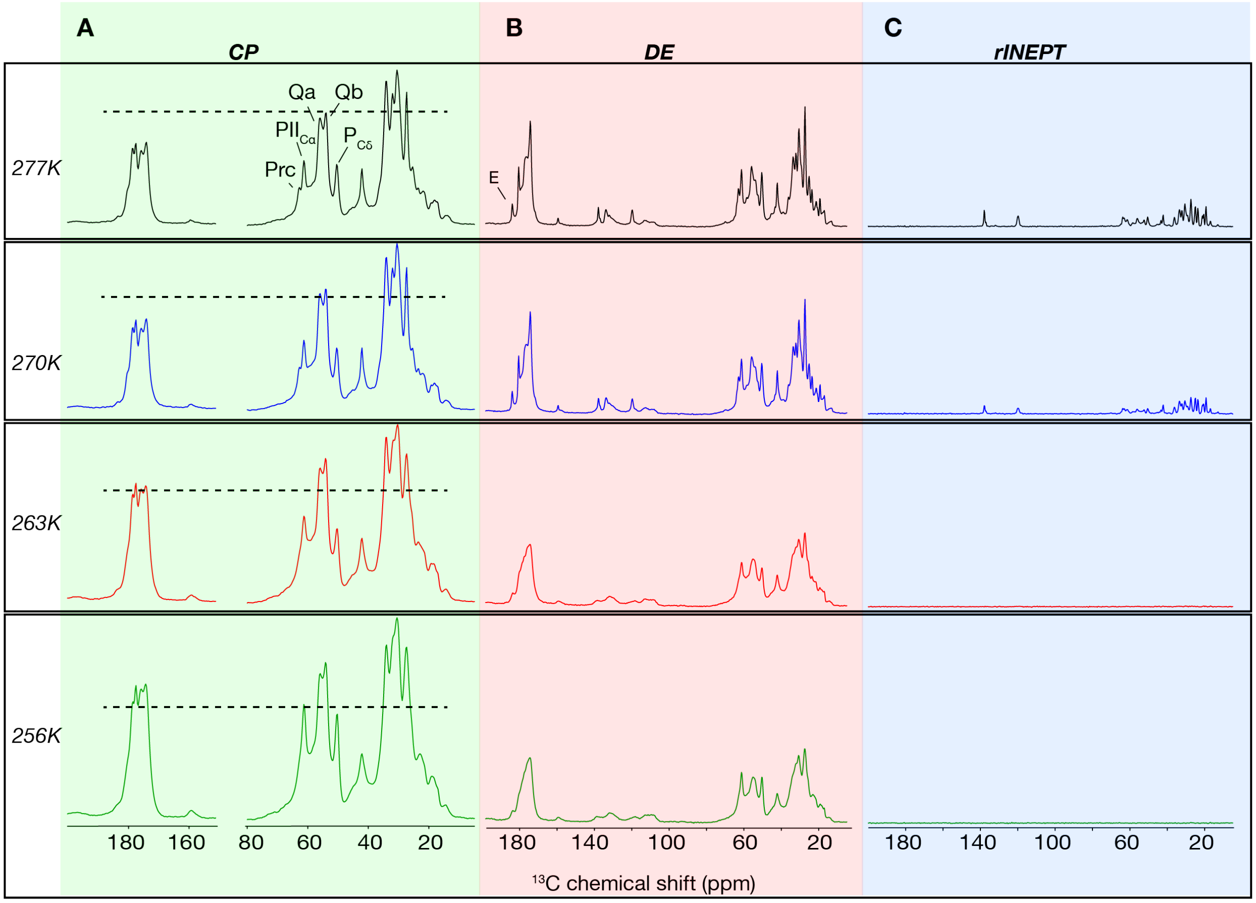


**Figure S1. Variable temperature 1D MAS ssNMR of Q44-HttEx1 fibrils.** (A) 1D $^{13}C$ CP-MAS ssNMR at 277, 270, 263 and 256K, reflecting signals of rigid or immobilized parts of the fibrils. (B) 1D $^{13}C$ direct excitation (DE) MAS ssNMR showing signals of both rigid and mobile residues. (C) 1D $^{13}C$ refocused INEPT showing highly mobile residues. All data acquired at 600 MHz ($^{1}H$) and 10kHz MAS rates. Assignments are based on ref. (Lin et al., 2017).

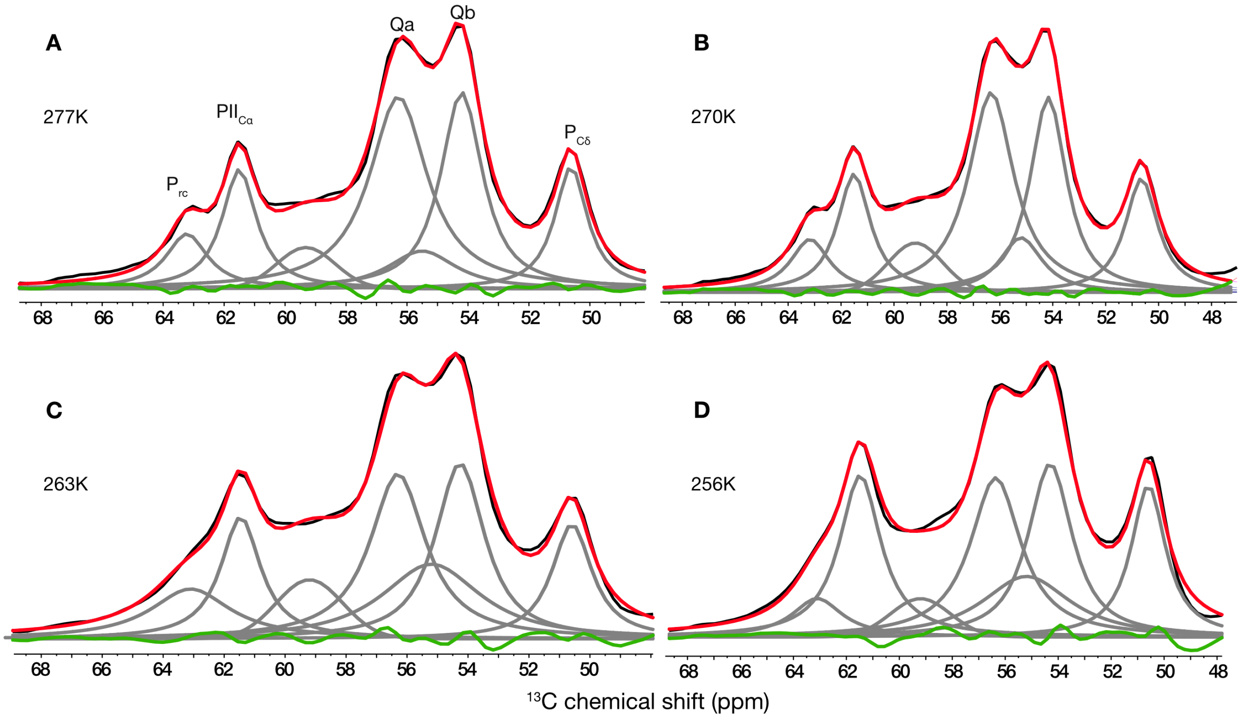


**Figure S2. Zoomed region of the 1D $^{13}$C CP spectra with deconvoluted glutamine and proline signals.** (A) 277K (B) 270K (C) 263K (D) 256K. Black line indicates experimental spectrum. Grey signals indicate deconvolution. Red spectrum is overall spectrum after deconvolution is applied. Green line indicates difference between experimental (black) and the spectrum after deconvolution (red).

**Table S2. Signal line widths of glutamine and proline peaks at different temperatures.** The uncertainty of linewidths is ±0.1 kHz. Signal areas are shown in arbitrary units. Abbrevations: Qa and Qb are the Cα peaks for the 'a' and 'b' conformers of the polyQ amyloid core; $PIIC_{\alpha}$ = Cα peak for polyproline II Pro residues; Prc = Cα peak for prolines in random coil; $PC_{\delta}$ = Cδ peak for all Pro residues ($PP_{II}$ and random coil overlapping).

| | **Qa** | | **Qb** | | **$PIIC_{\alpha}$** | | **$PC_{\delta}$** | | **Prc** | |
|---|---|---|---|---|---|---|---|---|---|---|
| **Temp. (K)** | **linewidth (kHz)** | **signal area** | **linewidth (kHz)** | **signal area** | **linewidth (kHz)** | **signal area** | **linewidth (kHz)** | **signal area** | **linewidth (kHz)** | **signal area** |
| 277 | 0.3 | $1.2\cdot10^{8}$ | 0.2 | $8.4\cdot10^{7}$ | 0.2 | $4.3\cdot10^{7}$ | 0.2 | $4.2\cdot10^{7}$ | 0.2 | $2.4\cdot10^{7}$ |
| 270 | 0.3 | $1.2\cdot10^{8}$ | 0.2 | $8.9\cdot10^{7}$ | 0.2 | $5.1\cdot10^{7}$ | 0.2 | $4.8\cdot10^{7}$ | 0.3 | $2.7\cdot10^{7}$ |
| 263 | 0.3 | $1.1\cdot10^{8}$ | 0.3 | $1.0\cdot10^{8}$ | 0.2 | $6.0\cdot10^{7}$ | 0.2 | $5.6\cdot10^{7}$ | 0.5 | $4.9\cdot10^{7}$ |
| 256 | 0.3 | $1.3\cdot10^{8}$ | 0.3 | $1.2\cdot10^{8}$ | 0.3 | $1.1\cdot10^{8}$ | 0.2 | $8.4\cdot10^{7}$ | 0.3 | $3.1\cdot10^{7}$ |

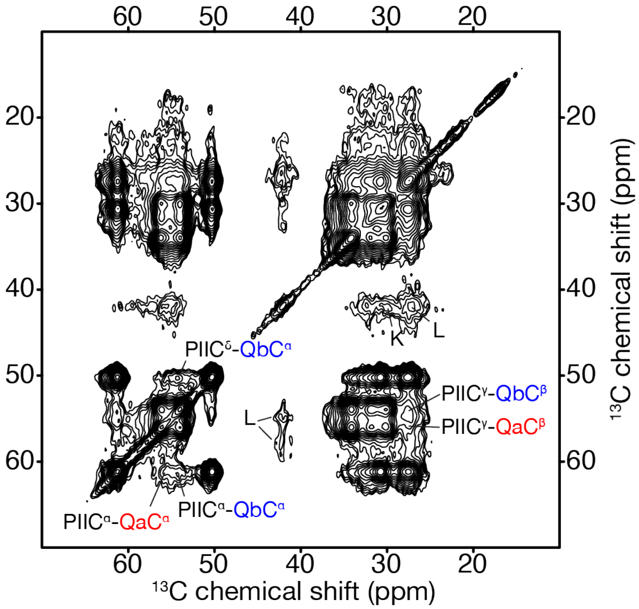


**Figure S3. 2D $^{13}C$-$^{13}C$ PDSD experiment on Q44-HttEx1 fibrils at 256K.** Long range magnetization transfer was performed using 500ms mixing time. Inter-residue correlations between prolines in PPII helix conformation in the C-terminus and type "a" and "b" glutamines are labeled.

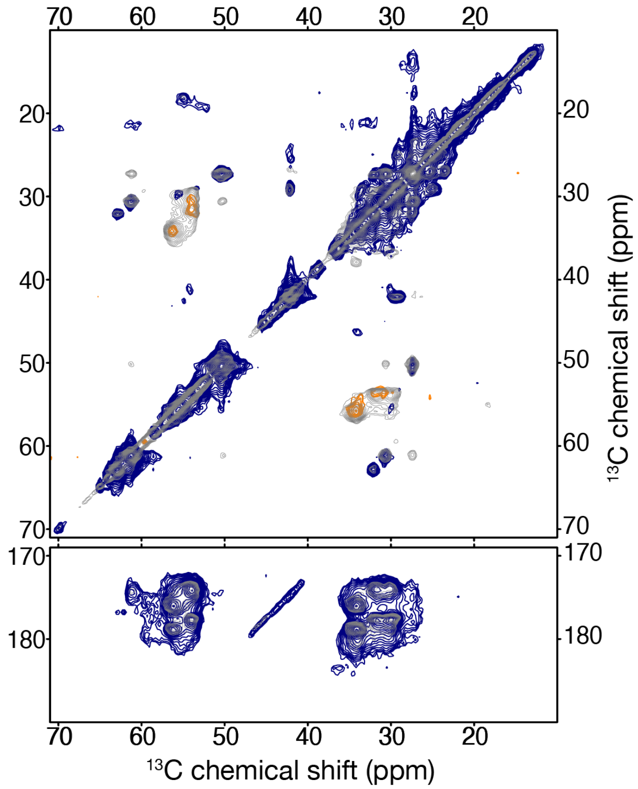


**Figure S4. Overlay of filtered and unfiltered 2D $^{13}C$-$^{13}C$ spectra of Q44-HttEx1 fibrils.** Grey spectrum represents the normal $^{13}C$-$^{13}C$ 2D DARR experiment (also shown in Figure 7A). The blue spectrum (with orange negative cross-peaks) shows the dynamically filtered $^{13}C$-$^{13}C$ DIPSHIFT-DARR 2D spectrum (Figure 7B). Both spectra were recorded at 277K with a DARR mixing time of 25ms.

**Table S3. Overview of selected $^{13}C$ ssNMR chemical shifts of glutamines of Q44-HttEx1 and other polyQ fibrils.** The uncertainty of chemical shifts is ±0.1-0.3ppm unless otherwise stated.

| | Chemical shifts (ppm) | | | | | Reference |
|---|---|---|---|---|---|---|
| **Residue** | **C'** | **$C_\alpha$** | **$C_\beta$** | **$C_\gamma$** | **$C_\delta$** | **-** |
| Gln "a" | 176.0 | 56.2 | 34.2 | 34.2 | 178.8 | - |
| Gln "b" | 174.0 | 54.0 | 31.7 | 30.0 | 177.7 | - |
| Gln "c" | - | 55.5 | 29.8 | 33.8 | 180.4 | - |
| Gln "d" | 173.6 | 53.3 | 29.0 | 29.0 | | - |
| Gln "c" published | - | - | - | 33.9 | 180.3 | (Hoop et al., 2016) |
| Q47c1 | 172.5 | 53.7 | 29.1 | 33.4 | 180.4 | (Hoop et al., 2014) |
| Q47c2 | 173.0+0.5 | 53.0 | 30.3 | 34.3 | 178.6 | (Hoop et al., 2014) |
| Q3 | 174.0 | 55.7 | 29.9 | 33.9 | 180.2 | (Schneider et al., 2011) |
| Gln "a" | 175.8 | 56.1 | 34.2 | 34.1 | 178.5 | (Isas et al., 2015) |
| Gln "b" | 173.9 | 54.0 | 31.8 | 29.9 | 177.4 | (Isas et al., 2015) |
| Gln "c" | 173.8 | 53.4 | 29.1 | 33.5 | 180.5 | (Isas et al., 2015) |
